# Enhanced RhoA signaling stabilizes E-cadherin in migrating epithelial monolayers

**DOI:** 10.1101/2021.04.08.439097

**Authors:** Shafali Gupta, Kinga Duszyc, Suzie Verma, Srikanth Budnar, Xuan Liang, Guillermo A. Gomez, Philippe Marcq, Ivar Noordstra, Alpha S. Yap

## Abstract

Epithelia migrate as physically coherent populations of cells. Earlier studies revealed that mechanical stress accumulates in these cellular layers as they move. These stresses are characteristically tensile in nature and have often been inferred to arise when moving cells pull upon the cell-cell adhesions that hold them together. We now report that epithelial tension at adherens junctions between migrating cells also reflects an increase in RhoA-mediated junctional contractility. We find that active RhoA levels were stimulated by p114 RhoGEF at the junctions between migrating MCF-7 monolayers, and this is accompanied by increased levels of actomyosin and mechanical tension. By applying a strategy to restore active RhoA specifically at adherens junctions by manipulating its scaffold, anillin, we found that this junctional RhoA signal was necessary to stabilize junctional E-cadherin during epithelial migration. We suggest that stabilization of E-cadherin by RhoA serves to increase cell-cell adhesion against the mechanical stresses of migration.

## Introduction

Cell migration is a complex mechanical process. Cells translocate by applying physical forces onto their environment, elements of the extracellular matrix (ECM) or other, adjacent cells. In turn, forces can be applied to migrating cells. This is exemplified by the collective migration of epithelia and endothelia, which characteristically move as coherent cell populations, whose constituent cells are physically linked together by cell-cell adhesions (Friedl and Gilmour, 2009; Friedl and Mayor, 2017). Studies using two-dimensional epithelial monolayers have shown that mechanical stresses build up within the cellular layer as they migrate (Reffay et al., 2014; Tambe et al., 2011; Trepat et al., 2009). These have been inferred to increase at cell-cell adhesions. As cells were typically crawling on ECM-based substrata via integrin adhesions in these experiments, the build-up in cellular stress has often been thought to reflect the transmission of traction forces through the cell to cell-cell junctions (Trepat and Fredberg, 2011). One simple model is that mechanical stresses at cell-cell adhesions would increase passively as migrating cells pull upon one another. Cadherin-based adherens junctions (AJ) are coupled to the F-actin cytoskeleton by intermediary proteins, such as α-catenin, vinculin and Myosin VI, that can bind directly to actin filaments (Lecuit and Yap, 2015). This physical coupling provides a way for traction forces to be transmitted through the actomyosin cytoskeleton to AJ.

But cadherin adhesions can also actively regulate their associated cytoskeleton, by recruiting actin regulators and signaling molecules that target the cytoskeleton (Lecuit and Yap, 2015). In particular, AJ in interphase epithelia are prominent sites of signaling by the RhoA GTPase. RhoA at AJ is activated by a number of upstream pathways, anchored by distinct guanine nucleotide exchange factors (GEFs) (Acharya et al., 2018; Garcia De Las Bayonas et al., 2019; Ratheesh et al., 2012) and RhoA signals to promote formin-based actin assembly and the activation of non-muscle myosin II (NMII) at AJ (Acharya et al., 2017; Cavanaugh et al., 2020; Curran et al., 2017; Ratheesh et al., 2012). In this report, we now show that RhoA signaling increases at AJ during epithelial migration, revealing that upregulation of active force-generation at AJ contributes to the build-up of tension at these cell-cell junctions. We further show that RhoA at AJ enhances E-cadherin stability during epithelial migration.

## Results and Discussion

### AJ contractility increases during epithelial migration

We performed our experiments using MCF-7 epithelial cells, a well-differentiated mammary cancer cell line that establishes robust AJ (Smutny et al., 2010). MCF-7 cells were grown to confluence in silicon moulds, which were then removed to allow the cells to migrate. Over the 12 h course of our experiments 12-15 rows of cells moved into the open space created by the mould. Lamellipodia were evident throughout the motile population, both at the leading margins of the sheet (Fig S1A) and also as cryptic lamellipodia within the moving sheet (Fig S1A, movie 1). Immunostaining for E-cadherin in 1-10 rows from migrating front confirmed that the cells had assembled AJ before the assays began and these were preserved in the migrating sheets (see Fig 4D).

Tensile stresses in migrating epithelia have often been characterized by inference techniques applied to traction force data (Tambe et al., 2011). To test if similar phenomena applied in our system, we performed traction force microscopy as MCF-7 cells migrated upon PDMS substrata and inferred stresses in the cellular layer by Bayesian Inversion Stress Microscopy (Nier et al., 2016; Ollech et al., 2020) (see Methods for details). Stresses were predominantly tensile in the pre-migratory MCF-7 monolayers, indicating that the cells were not over-confluent or compressed before the start of the experiment (Figure 1A). After the mould was removed, these tensile stresses increased with migration (Fig 1B). Thus, migrating MCF-7 cells showed increases in monolayer tension similar to what has earlier been reported for other types of epithelial cells.

**Figure 1.**
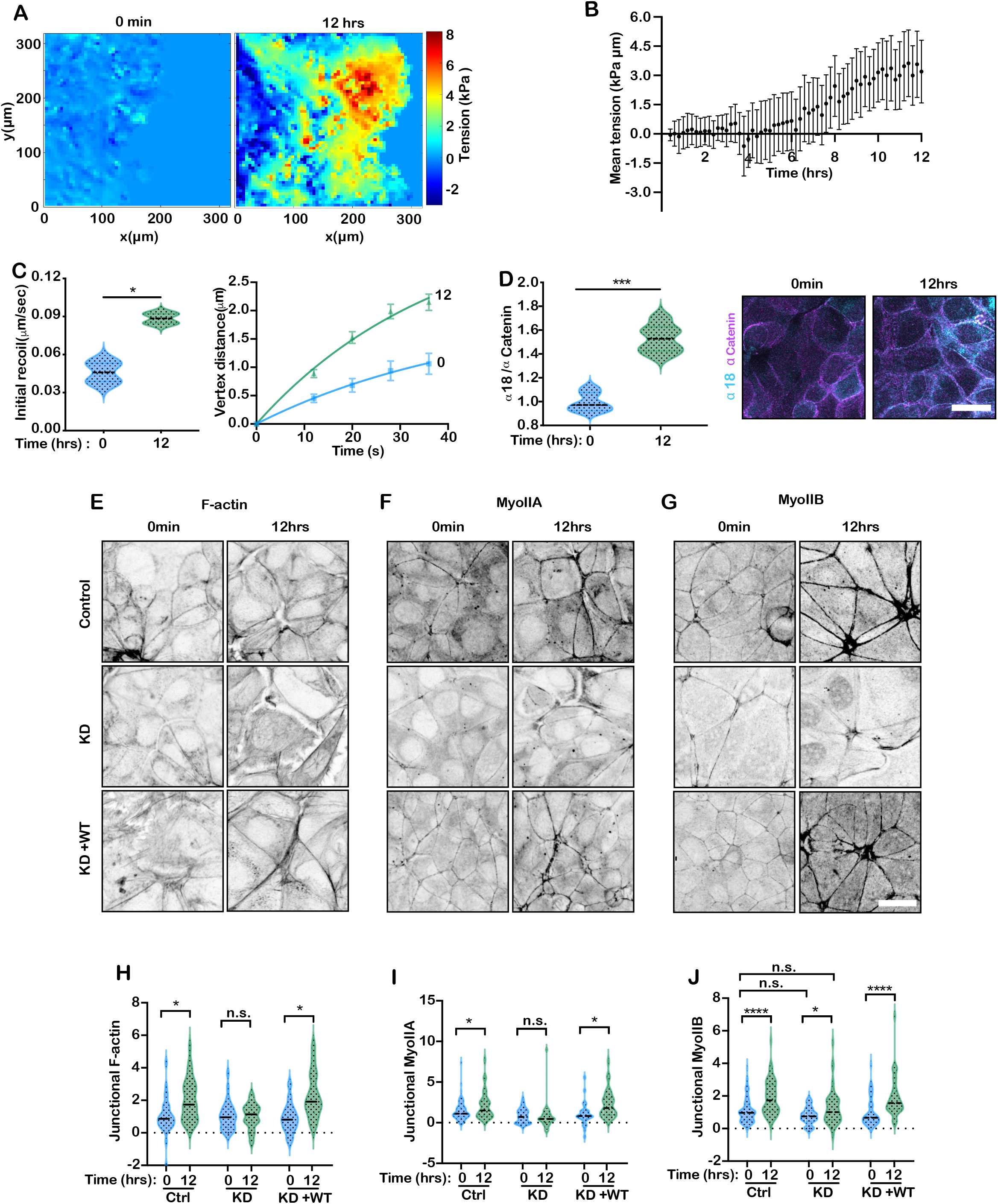
AJ tension increases in migrating epithelial cells. (A,B) Stress in Caco-2 epithelial monolayers before (0 hr) and for 12 h during migration calculated by Bayesian Inversion Stress Microscopy. Representative tension maps (A) and quantification of mean tension (B). (C) AJ tension inferred from recoil velocity after laser ablation of AJ. Initial recoil velocities (left) and recoil curves (right). (D) Tension at the cadherin-catenin complex inferred from immunostaining for α18 mAb at AJ (normalized to α-catenin staining): quantification (left) and representative images (right). (E-J) Junctional F-actin (E,H), Myosin IIA (F,I) and Myosin IIB (GI) in migrating MCF-7 cells expressing control shRNA, Anillin shRNA (KD), or Anillin shRNA reconstituted with full-length GFP-Anillin (KD+WT). Representative images (E-G) and quantification (H-J). Data are means ± SEM; n=3 independent experiments. n.s., not significant; *,P<0.05; **,P<0.01; ***,P<0.001; ****,P<0.0009; unpaired t-test (C,D) or one-way ANOVA (H,I,J) with Dunnett’s multiple comparisons test. Scale bars: 20 μm.

To more specifically evaluate changes in junctional tension in migrating cells, we identified AJ by expressing E-cadherin-mCherry and measured their initial recoil after being cut by laser ablation (Ratheesh et al., 2012). As shown in Fig 1C, a basal recoil was evident at AJ in premigratory monolayers and this increased with migration, consistent with an increase in tension within the AJ between migrating cells. To assess whether molecular-level tension changed at the AJ, we then immunostained cells for the α-18 epitope of α-catenin, which is a cryptic site that is revealed when tension is applied to the α-catenin molecule (Yonemura et al., 2010). MAb α-18 staining, expressed as a ratio of that for α-catenin itself, increased in migrating monolayers (Fig 1D). Together, these findings imply that tension increases at AJ of migrating MCF-7 cells.

This observed increase in AJ tension could have been occurred if AJ passively resisted forces of migration being transmitted from elsewhere in the cells and/or if contractility at AJ was itself enhanced. We examined this further by evaluating actomyosin at AJ. Phalloidin staining revealed that levels of cortical F-actin increased at the cell-cell contacts (Fig 1E,H). MCF-7 cells express all three mammalian NMII paralogs (Smutny et al., 2010) and antibodies were available that effectively recognized NMIIA and NMIIB(Fig 1F,G). Indeed, quantitative immunofluorescence showed that both NMIIA and NMIIB increased at AJ in migrating cells (Fig I,J). Overall, this increase in actomyosin suggested that upregulation of cortical contractility may have contributed to increasing tension at AJ during collective migration.

### RhoA signaling is upregulated at AJ during collective epithelial migration

To explore whether contractility was actively upregulated at AJ between migrating cells we focused on the RhoA GTPase, a canonical regulator of cellular actomyosin that is also active at AJ (Priya et al., 2013; Ratheesh et al., 2012). We therefore investigated whether junctional RhoA is also altered during MCF-7 sheet migration.

To identify the sites of RhoA signaling in collectively-migrating MCF-7 cells, we expressed AHPH, a location biosensor that identifies active, GTP-loaded RhoA (Piekny and Glotzer, 2008; Priya et al., 2015). As previously observed (Priya et al., 2015), AJ were prominent sites of GTP-RhoA in premigratory monolayers (Fig 2A; note however that image intensity in the images presented was reduced to avoid saturation, Fig S2A). Strikingly, AHPH levels at junctions increased significantly when cells began to move (Fig 2A; Movie2). This increase was most pronounced in the first 5 rows of cells, but became evident in the cells further behind as they began to migrate (Fig 2A,B). AHPH was also evident in the free lamellipodia of the leader cells (Fig S2B), but this did not increase with time, in contrast to the junctional signal (Fig 2C). The increase in junctional RhoA signaling was confirmed using a FRET-based RhoA activity sensor (Fig S2C). Together, these findings indicated that RhoA signaling is upregulated at AJ when MCF-7 cells migrate.

**Figure 2.**
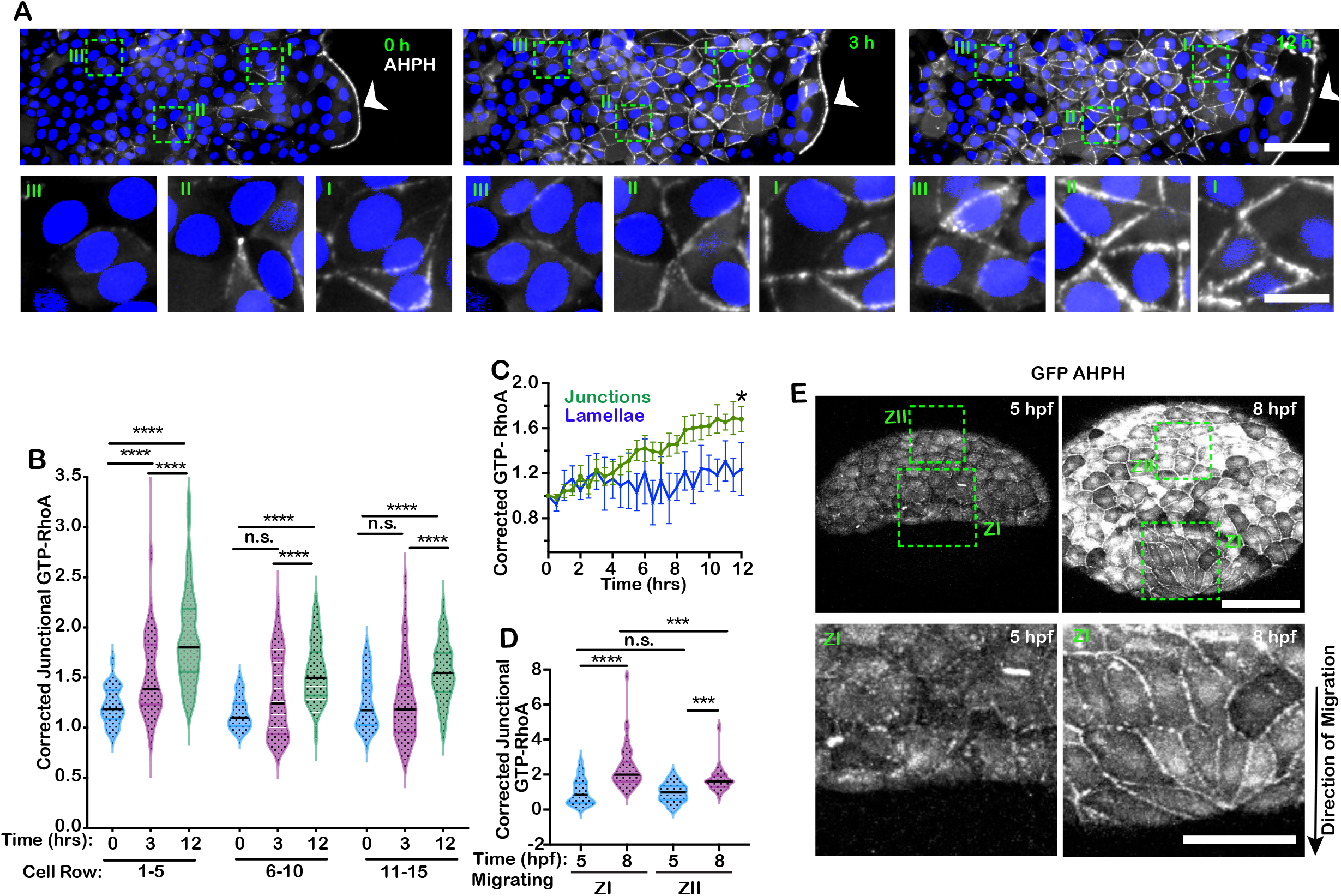
RhoA signaling is upregulated at the AJ of migrating epithelial cells. GTP-RhoA was detected using the AHPH location biosensor (GFP-AHPH_ in MCF-7 cells (A-C) and zebrafish embryos (D,E). (A) Representative images of GFP-AHPH in migrating MCF-7 monolayers. Dashed boxes show different zones of migrating monolayer (I = cell row 1-5, II = cell row 6-10, III = cell row 11-15). (B) Quantification of junctional GFP-AHPH (corrected for cytosolic expression in each cell) at different zones of the migrating monolayer. (C) Change in junctional GFP-AHPH compared with GFP-AHPH at the lamellae of leader cells. (D, E) GFP-AHPH during epiboly of in *tg(krt4:GFP-AHPH)*^uqay2^ zebrafish embryos at 5 and 8 hour post fertilization (hpf): quantification of junctional GFP-AHPH (D, corrected for cytosolic expression) and representative images (E). Dashed boxes show different zones of migrating monolayer. Data are means ± SEM; n=3 independent experiments. n.s., not significant; *,P<0.05; **,P<0.01; ***,P<0.001; ****,P<0.0009; one-way ANOVA (B,C,D) with Dunnett’s multiple comparisons test. Scale bars: A, top panel: 100μm; A, bottom panel: 20μm; E: 25μm.

To test this observation in another system, we used a transgenic zebrafish line expressing GFP-AHPH [*tg(krt4:GFP-AHPH)*^uqay2^] (Duszyc et al., 2021) and studied its behaviour during epiboly, a form of collective migration in the early vertebrate embryo (Bruce and Heisenberg, 2020). *tg(krt4:GFP-AHPH)*^uqay2^ embryos showed AHPH staining at their medial-apical cortex as well as at cell-cell junctions. During early epiboly AHPH was first evident in puncta at the medial-apical surface of the cells, then became prominent at junctions by 50% epiboly, where it increased as epiboly progressed (Fig 2D,E; Movie 3). Therefore, the upregulation of junctional RhoA signaling may be a general feature of collective epithelial migration. This increase in GTP-RhoA provided a plausible candidate to enhance contractile tension at AJ.

### p114 RhoGEF participates in activating junctional RhoA during migration

To gain insight into the mechanism responsible for upregulating junctional RhoA, we sought to identify its upstream activator. A number of GEFs have been reported to stimulate RhoA at AJ (Garcia De Las Bayonas et al., 2019). In mammalian epithelial cells, these include Ect2, which appears to support steady-state RhoA signaling at interphase AJ (Ratheesh et al., 2012), and p114 RhoGEF, which can increase junctional RhoA activity when tensile forces are applied to epithelia (Acharya et al., 2018). We focused on p114 RhoGEF, which showed faint immunostaining at AJ in premigratory MCF-7 monolayers, and this increased with migration (Fig 3A,B). Total cellular levels of p114 RhoGEF did not change (Fig 3C), suggesting that the increased junctional staining was due to enhanced recruitment.

**Figure 3.**
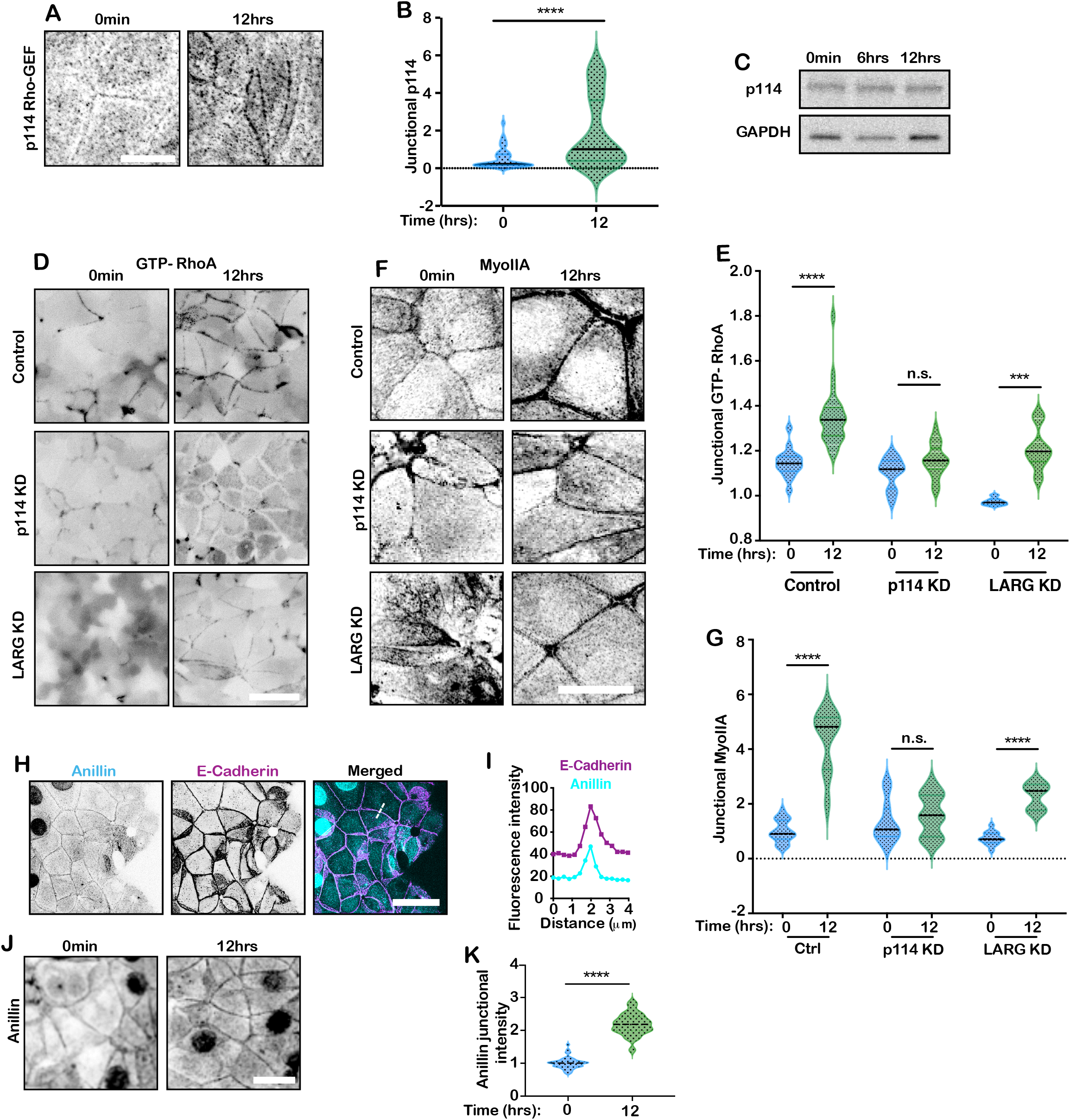
p114 RhoGEF and anillin are AJ components during epithelial migration. (A,B) Junctional p114 RhoGEF in migrating MCF-7 monolayers: representative images (A) and quantification (B). (C) Representative immunoblots for total p114 RhoGEF in migrating MCF-7 monolayers (0 – 12 hr). (D-G) Effect of p114 RhoGEF RNAi on RhoA signaling at AJ in migrating MCF-7 monolayers. Monolayers were transfected with control siRNA (control), p114 RhoGEF RNAi (p114 KD) and LARG RNAi (LARG KD). Representative images and quantitation for GFP-RhoA (detecting GTP-RhoA; D, E) and Myosin IIA (F,G). (H-I) Colocalization of anillin with E-cadherin in a representative migrating monolayer: immunofluorescence images and quantitation. (J-K) Anillin increases AJ in migrating MCF-7 monolayers: representative images (J) and quantification (K). Data are means ± SEM; n=3 independent experiments. n.s., not significant; *,P<0.05; **,P<0.01; ***,P<0.001; ****,P<0.0009; unpaired t-test. Scale bars: A,E,G,J: 20μm; D: 50μm.

Then we asked if p114 RhoGEF RNAi (S3A) affected RhoA signaling at AJ during epithelial migration. p114 RhoGEF depletion did not alter baseline levels of junctional GTP-RhoA, but it prevented GTP-RhoA from increasing when the cells migrated (Fig 3D,E). Consistent with this, junctional NMIIA levels did not increase when p114 RhoGEF KD cells migrated (Fig 3F,G). In contrast, depleting LARG, another mechanically-activated RhoGEF (Guilluy et al., 2011), did not prevent GTP-RhoA from increasing at the AJ during migration (Fig 3D,E). This implied that p114 RhoGEF was critically responsible for enhancing RhoA at AJ when MCF-7 cells migrated. As p114 RhoGEF can activate RhoA when tension is applied to AJ (Acharya et al., 2018), it is possible that during epithelial migration it responds to tugging forces that moving cells apply to one another. p114 RhoGEF would then provide a mechanism for active junctional contractility to be elicited when traction forces are transmitted to AJ.

### Anillin depletion prevents the upregulation of junctional RhoA

Our end-goal was to understand how RhoA might affect AJ function in migrating cells. However, manipulation of p114 RhoGEF presented interpretative complexities, because this GEF also activates RhoA at other sites within cells (Kelly et al., 2007), including tight junctions between epithelial cells (Terry et al., 2011). To focus on the impact of RhoA at AJ, we therefore required an alternative approach that would allow us to manipulate it specifically at these junctions.

For this purpose, we exploited our recent discovery that the cytoskeletal protein, anillin, is a kinetic scaffold for GTP-RhoA, which stabilizes it and promotes its cortical signaling (Budnar et al., 2019). Anillin localizes to AJ (Fig 3H,I) (Budnar et al., 2019) and, indeed, increased at AJ in migrating cells (Fig 3J,K). Depleting anillin by RNAi (knock-down, KD, Fig S3C) reduced junctional GTP-RhoA in premigratory monolayers, as we had observed earlier (Budnar et al., 2019), and this was restored by expression of an RNAi-resistant transgene (Fig S3C). It should be noted that although AHPH derives from the C-terminus of anillin, it is not affected by manipulating levels of anillin (Budnar et al., 2019).

Strikingly, although anillin KD cells retained contact with one another as they migrated (Fig 4D), they did not upregulate GTP-RhoA at their junctions (Fig 4B,C). Consistent with this, anillin KD also specifically prevented NMII levels from increasing at the AJ of migrating cells (Fig 1E,G). This suggested that anillin was necessary for junctional RhoA to be upregulated during epithelial migration. However, anillin is found at other parts of the cell cortex and, indeed, anillin KD also reduced GTP-RhoA levels in the lamellae of leader cells (Fig S2B).

**Figure 4.**
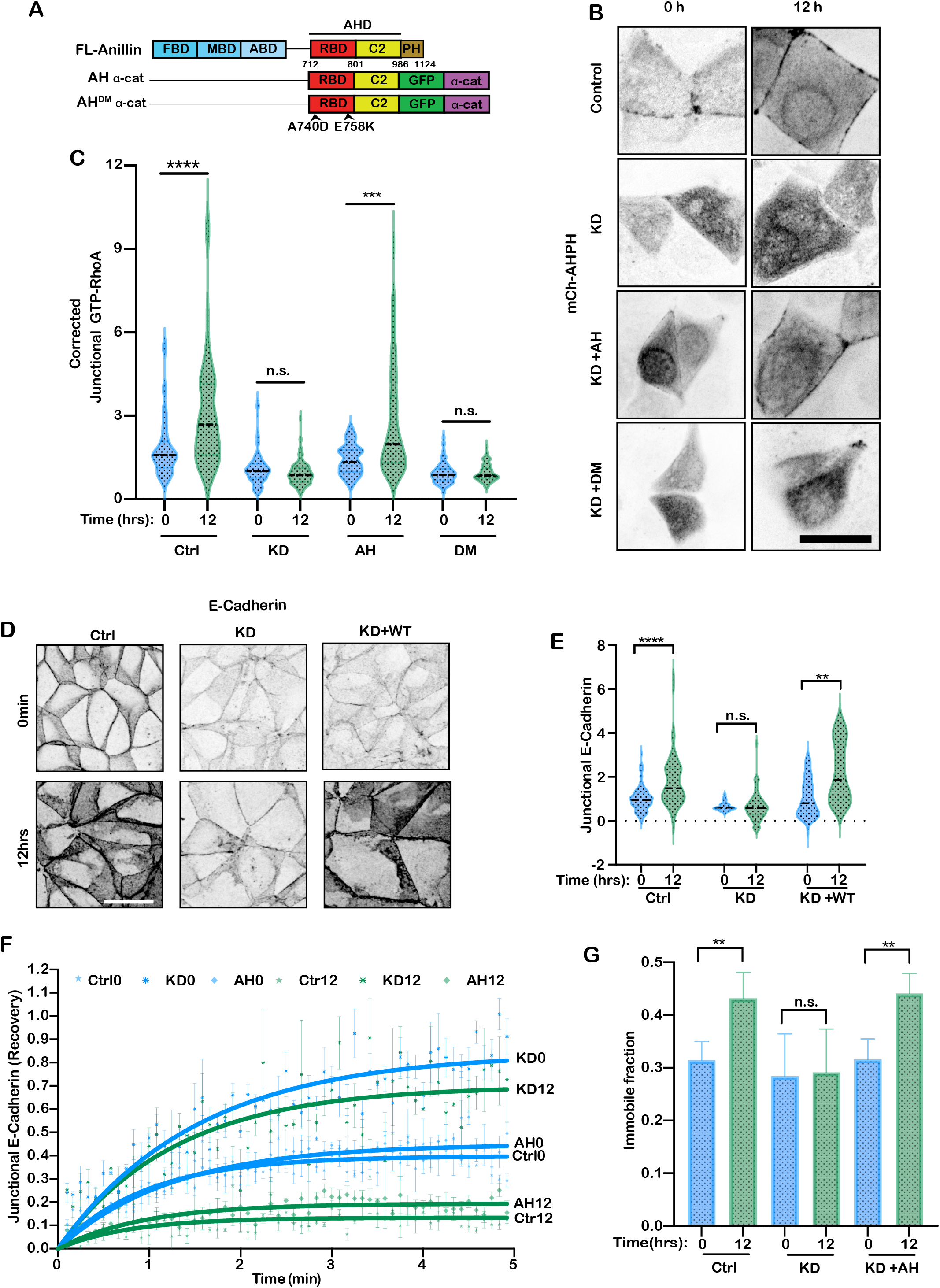
Junctional RhoA stabilizes E-cadherin in migrating monolayers. RhoA was selectively restored at AJ by targetting the GTP-RhoA-binding domain of anillin (AHPH) to E-cadherin in anillin RNAi cells. (A) Cartoon depicting the structure of the strategy. AHPH or its RhoA-uncoupled (DM) mutant were fused to the N-terminus of α-catenin, as described in detail previously (Budnar et al., 2019). (B,C) Effect on junctional GTP-RhoA (mCherry-AHPH) of expressing AH-α-catenin (AH) or AH^DM^-α-catenin (DM) in anillin KD cells. Representative images (B) and quantification (C). AH-α-catenin, but not AH^DM^-α-catenin, restores junctional GTP-RhoA levels to anillin KD cells. (D,E) Impact of junctional RhoA signaling on E-cadherin levels at AJ in migrating MCF-7 monolayers: representative immunofluorescence images (D) and quantification (E). (F,G) Effect of junctional RhoA signaling on the stability of junctional E-cadherin, measured by fluorescence recovery of transiently-expressed E-cadherin-mCherry. FRAP curves (F) and immobile fractions (G). Data are means ± SEM; n=3 independent experiments. n.s., not significant; *,P<0.05; **,P<0.01; ***,P<0.001; ****,P<0.0009; unpaired t-test. Scale bar: 20μm.

Therefore, to selectively manipulate RhoA at AJ, we reconstituted anillin KD cells with a chimeric protein consisting of the anillin homology (AH) domain that was targeted to AJ by fusion with α-catenin (AH-α-catenin; (Budnar et al., 2019)). The AH domain is responsible for kinetic scaffolding of GTP-RhoA, so we reasoned that this strategy would restore its activity selectively at AJ (Fig 4A,S4A). Indeed, expression of AH-α-catenin restored the capacity for migrating KD cells to upregulate GTP-RhoA at their junctions, despite anillin being otherwise depleted in the cells (Fig 4B,C). However, the upregulation of junctional GTP-RhoA was not restored by an AH^DM^-α-catenin mutant that cannot bind and stabilize GTP-RhoA (Fig 4B,C; (Budnar et al., 2019). Therefore, AH-α-catenin could selectively restore RhoA signaling to junctions after anillin KD. Combining anillin KD with AH-α-catenin reconstitution then provided a strategy to selectively interrogate the impact of junctional RhoA signaling on AJ in epithelial migration.

### Upregulation of RhoA stabilizes E-cadherin in migrating monolayers

We then used this strategy to test how upregulation of RhoA at AJ might affect E-cadherin. As RhoA signaling stabilizes E-cadherin at AJ (Priya et al., 2013; Ratheesh et al., 2012), we asked if the increased junctional RhoA observed in migrating MCF-7 cells might affect E-cadherin at their junctions. Indeed, immunostaining showed that E-cadherin levels increased in migrating cells. Furthermore, this was reduced by anillin RNAi and selectively restored by expression of an RNAi resistant anillin transgene (Fig 4D,E).

Then, we used fluorescence recovery after photobleaching (FRAP) assays to measure the turnover of E-cadherin-mCherry. Control cells stabilized E-cadherin-mCherry as they migrated, as reflected in an increased immobile fraction (~ 27% at 0 h to ~ 39% at 12 h). Strikingly, E-cadherin was more dynamic in premigratory Anillin KD cells (immobile fraction ~ 12%) and this failed to become more stable upon migration (immobile fraction ~ 18% at 12 h). Importantly, stabilization of E-cadherin during migration was restored to anillin KD cells by expression of AH-α-catenin. The premigratory immobile fraction was increased, compared with KD cells and stabilized further upon migration (Fig 4F,G). This indicated that E-cadherin was stabilized as a specific consequence of RhoA signaling at the AJ. Together, these findings indicated that the enhanced junctional RhoA signaling seen in migrating MCF-7 monolayers enhanced the stability of E-cadherin in these collectively-moving cells.

## Conclusion

Together, our results show that RhoA is activated by p114 RhoGEF at the AJ of migrating MCF-7 epithelial monolayers. This is accompanied by an increase in cortical actomyosin at those junctions, suggesting that tensile stress increases at the AJ of migrating cells through local up-regulation of RhoA, as well as any forces that applied to junctions when moving cells pull on one another. Indeed, the involvement of p114 RhoGEF in mechanosensitive RhoA activation (Acharya et al., 2018) may provide a way for passively applied forces to elicit an enhanced active contractile response at AJ. In other words, the aggregate increase in tensile stresses at the junctions of migrating epithelia may reflect a combination of traction forces transmitted to junctions and local responses to those transmitted forces.

We further find that junctional RhoA enhances the stability of E-cadherin at the junctions between migrating cells. Potentially, this could serve to reinforce AJ against pulling forces that they experience during migration (Acharya et al., 2018). A fundamental role of cell-cell adhesion is to resist detachment forces that can separate cells from one another. Accordingly, a variety of mechanisms exist that can enhance cadherin adhesion when disruptive mechanical forces are applied (Charras and Yap, 2018; Rakshit et al., 2012). These include catch bonds found in the adhesive binding interaction of the extracellular domain (Rakshit et al., 2012) and also in the association of α-catenin to actin filaments (Buckley et al., 2014; Ishiyama et al., 2018). In an earlier study, we found that tension-activated RhoA signaling can protect epithelia by increasing the stiffness of the junctional cytoskeleton (Acharya et al., 2018). The stabilization of E-cadherin that we have now observed would provide an additional line of protection by increasing the levels of this adhesion receptor present at the AJ, thereby increasing the adhesion forces that resist detachment.

## Acknowledgements

We thank our lab colleagues for their support and advice during this project. This work was funded by the NHMRC Australia (GNT 1123816 and 1140090, and Fellowship 1136592 to AY) and Australian Research Council (DP19010287 and DP190102230 to ASY, FT160100366 to GAG). IN was supported by the European Molecular Biology Organization (EMBO ALTF 251-2018). Collaboration between PM and ASY was supported in part by the National Science Foundation under Grant No. NSF PHY-1748958. Optical microscopy was performed at the ACRF/IMB Cancer Biology Imaging Facility, established with the generous support of the Australian Cancer Research Foundation.

## Materials and Methods

### Cell culture and transfection

MCF-7 cells were maintained in Dulbecco’s Modified Eagle Medium (DMEM, Gibco) supplemented with 10% FBS, 1% NEAA, 1% L-glutamine (Gibco) and pen-strep (Gibco). HEK-293T cells were maintained in DMEM supplemented with 10% FBS-HI, 1% NEAA, 1% L-glutamine and pen-strep. Cell lines were maintained in 25 μg/ml plasmocin (Invitrogen) to prevent mycoplasma contamination and were routinely screened for the presence of mycoplasma. Barrier-separated cell monolayers were generated by seeding MCF-7 cells at a confluency of 1.5 × 10^5^ cells/ml in two-well culture insertion barriers (ibidi), pre-placed in glass bottom dishes (Shengyou Biotechnology). After 100% confluency was reached, the barriers were removed, and monolayer migration was monitored. For live-cell imaging, culture medium was replaced by Hank’s balanced salt solution supplemented with 10% FBS, 10 mM HEPES (pH 7.4) and 5 mM CaCl_2_. Transient transfections were performed after seeding using Lipofectamine 2000 or 3000 for DNA (Invitrogen) and RNAiMAX for RNA (Invitrogen). Mixtures were prepared according to manufacturer’s protocol.

### Zebrafish husbandry

Adult Zebrafish and embryos were maintained by standard protocols approved by the University of Queensland Animal Ethics Committee.

### Live imaging of epiboly migration

Generation of the *tg(krt4:GFP-AHPH*)^uqay2^ line was described recently (Duszyc et al., 2021). *Tg(krt4:GFP-AHPH*)^uqay2^ fish were crossed to wild-type fish. Heterozygous transgenic 4.5hpf embryos were dechorionated using forceps and mounted in low-melting agarose (1% in E3 embryo medium) in glass-bottom imaging dishes.

### Lentiviral infection

Lentivirus packaging was performed by Lipofectamine 2000-based co-transfection of HEK293T cells with packaging vectors (VSV-G, RSV-rev, pRRE) and the lentiviral vector PLL5.0 containing the genes of interest in OptiMEM. 8 hours after transfection the media was replaced by normal culture media. 10 mM sodium butyrate and fresh culture media was added 24 hours after transfection to reduce any pH changes. The virus-containing supernatant was subsequently harvested by centrifugation and concentrated on Millipore spin columns. Virus was stored at −80°C or used to infect target cells.

To infect target cells, concentrated virus was mixed with 8 μM polybrene and added to the cells. 24 hours after infection, the medium was replaced by normal culture medium. The following constructs were expressed through lentiviral infections: GFP-AHPH (gift from M. Glotzer, University of Chicago, USA), GFP alone, Anillin KD, Anillin KD+R, Anillin KD-AH GFP α-Catenin, Anillin KD-DM GFP α-Catenin.

### Immunofluorescence staining

For immunofluorescence staining, cells were cultured on glass coverslips and fixed with 4% PFA (α-catenin, α18, MRLC, pMRLC), 10% TCA (RhoA) or −20°C Methanol (Anillin, Myosin IIA, Myosin IIB, Actin, E-Cadherin, p114 RhoGEF) for 15 minutes. TCA fixation was terminated with 30 mM glycine in PBS for 5 minutes. After fixation, cells were permeabilised with 0.25% Triton X-100 for 5 minutes. Next, samples were blocked with 5% fat-free dry milk in PBS and subsequently incubated with primary antibodies overnight at 4°C and fluorescently labelled secondary antibodies for 2 hours at RT. Finally, samples were washed and mounted in ProLong Gold (Cell Signaling Technology).

### Western blotting

For western blot, cells were lysed in 1x sample buffer (4x: 200mM Tris-Cl, 40% glycerol, 8% SDS and 0,4% BPB) and incubated for 10 min at 100°C. Samples were run on 4, 8 or 15% SDS polyacrylamide electrophoresis gels (depending on the molecular weight of the proteins of interest) and transferred onto nitrocellulose membranes. The nitrocellulose membranes were subsequently blocked with 5% non-fat milk for 1 hour, incubated with primary antibodies overnight at 4°C and incubated with species-specific horseradish peroxidase-conjugated secondary antibodies for 1 hour. Next, the membranes were developed using enhanced chemiluminescence (ECL).

### Antibodies and chemicals

For immunofluorescence, mouse antibodies against the following proteins were used: MRLC (Abcam ab594; 1:100), Anillin (Santa Cruz SC-271814; 1:100), Myosin IIB (Abcam ab684; 1:300), Actin (Thermo Fisher Scientific A5228; 1:100) and RhoA (Santacruz SC-418; 1:100). Rabbit antibodies against the following proteins were used: α-catenin (Thermo Fisher Scientific 711200; 1:100), pMRLC (Merck Millipore AB3381; 1:50) and Myosin IIA (Biolegend 909801; 1:300). Rat antibodies against the following proteins were used: α18 (gift from Professor A. Nagafuchi, Nara Medical University, Japan; 1:50) and rat E-Cadherin (Abcam 13-1800; 1:500). Goat antibody against p114 RhoGEF was purchased from Sapphire Bioscience (EB06163; 1:100). AlexaFluor 488-, AlexaFluor 547- and AlexaFluor 647-conjugated antibodies against mouse, rabbit, rat and goat were purchased from Thermo Fisher Scientific (1:500). AlexaFluor 647-conjugated phalloidin was purchased from Thermo Fisher Scientific (A22287).

For western blot, mouse antibodies against the following proteins were used: Anillin (Santa Cruz SC-271814; 1:200), GFP (Sigma-Aldrich 11874460001; 1:2000). Rabbit antibodies against the following proteins were used: GAPDH (Trevigen 2275-PC-100; 1:5000), p114 RhoGEF (Abcam ab96520; 1:100), Alpha-catenin (Thermo Fisher Scientific 711200; 1:500). HRP-conjugated antibodies against mouse and rabbit were purchased from Bio-Rad Laboratories (1:5000)

### DNA constructs and siRNAs

The following DNA constructs were used in this study: pLL5.0-GFP, pLL5.0-Anillin shRNA and pLL5.0-Anillin-shRNA-GFP-Anillin FL have been published pBudnar *et al.*, 2019). PLL5.0 E-cadherin-mCherry KDR has been pubished (Wu *et al.*, 2015). pLL5.0-Anillin ShRNA-CMV-AHPH-GFP-⍰-cat FL and PLL5.0-Anillin ShRNA-CMV-AHPH^DM^-GFP-α-cat FL were cloned for this study. DNA maps are available upon request.

The following siRNAs were used for transient gene silencing: Anillin: 5’- ATGCCTCTTTGAATAAATT-3’ (Sense) + 5’-TTTATTCAAAGAGGCATCG-3’ (Antisense), P114RhoGEF a: 5’-TCAGGCGCTTGAAAGATATT-3’ (Sense) + 5’-TATCTTTCAAGCGCCTGATG-3’ (Antisense), P114RhoGEF b: 5’- GGACGCAACTCGGACCAATTT-3’ (Sense) + 5’- ATTGGTCCGAGTTGCGTCCCA-3’ (Antisense), LARG a: 5’- GGCAACATTTCCCAAGATATT-3’ (Sense) + 5’-TATCTTGGGAAATGTTGCCAG-3’ (Antisense), LARG b: 5’- GATCAAATCTCGTCAGAAATT-3’ (Sense) + 5’-TTTCTGACGAGATTTGATCAT-3’ (Antisense). All siRNAs were purchased from Sigma-Aldrich.

### Traction Force Microscopy sample preparation

Polyacrylamide gels with a Young’s modulus of 15 kPa were prepared using CY-52-276 A and CY-52-276 B (Dow Corning Toray) in 1:1 v/v ratio. Gels were spin coated at 500 RCF for 30 secs to get an even coating and cured for 2 hours at 80°C. 5% APTES (Sigma-Aldrich A3648) in ethanol was added to the gels followed by an incubation of 10 minutes. The solution was then removed, and the gels were washed with ethanol and dried at 80°C for 30 minutes. 200 nm nanobeads (Thermo Fisher Scientific, FluoSpheres®) were diluted at 1:30000 with distilled water and sonicated for 10 minutes to break the clumps. The silane-PDMS was functionalised with the nanobeads solution for 5 minutes at room temperature after which the beads were removed again. Functionalised gels were rinsed with distilled water and dried for 15 minutes at 80°C. Beads were then passivated with 100 mM Tris for 10 min at RT. Subsequently, the solution was removed, gels were washed with distilled water and dried in the oven for 15 minutes at 80°C. 10 ⍰g/ml fibronectin was added to the gels followed by 1 hour incubation at room temperature. Next, the gels were incubated with 1% Pluronic for 30 mins to sterilise and washed with PBS. Finally, cells were added and monitored until a confluent monolayer was generated.

### Microscopy and image analysis

The following microscopes were used in this study: 1) Traction Force Microscopy experiments were performed on a Nikon Ti-E deconvolutional microscope (20X, 0.75 NA Plan Apo objective) driven by NIS –Elements AR software. 2) Laser ablation experiments were performed on a LSM 510 meta Zeiss confocal microscope (63X, 1.4 NA Plan Apo objective) driven by Zen black software. 3) FRAP experiments were performed on an inverted Zeiss Meta LSM 510 Zeiss confocal microscope (100X, 1.4 NA Plan Apo objective) driven by Zen black software. 4) FRET experiments were performed on a LSM 710 Meta confocal (63X 1.4 NA Plan Apo objective) with specific detector for CFP/YFP imaging. Microscope was driven by Zen black software. 5) Immunofluorescence images were acquired with an inverted Zeiss 880 or an upright 710 Meta laser-scanning microscope (63X, 1.4 NA Plan Apo objective or 100X, 1.4 NA Plan Apo objective) driven by Zen black software. 6) Zebrafish were imaged on a Zeiss LSM710 confocal microscope (20×, 0.8NA Plan Apo objective).

### Traction Force Microscopy and stress analysis

Cells were observed by using bright-field microscopy. Nanobeads were observed using the far-red channel. At the end of the experiment, cells were trypsinised to get null force. The gel deformations were obtained by recording the displacement of nanobeads in comparison to the null force. PIV lab toolbox (MATLAB) was used to obtain velocity vectors by saving images in the sequence (1-2,3-4,5-6,..) and not (1-2,2-3,3-4,…). When acquiring movies using a 20X objective, 3 Pass filters were used, 128(64), 64(32), 32(16). After PIV, outliers were removed manually. Vector validation was conducted by defining, and then refining, velocity limits. BISM (Bayesian Inversion Stress Microscopy) was run to back calculate the internal stress field of a cell monolayer from the traction forces data. For this, we analysed regions of 300 μm x 300 μm, applying stress-free boundary conditions only on the cell-free boundary (on the right-hand-side of Fig1A). As in (Ollech et al., 2020) we validated BISM used with these boundary conditions by comparing tension values obtained by spatially-averaging the 2D tension field over the direction parallel to the migrating front, with the 1D tension obtained from direct integration of traction forces along the direction perpendicular to the migrating front (Trepat et al., 2009) (not shown). Images in Fig1A,B show isotropic stress (or tension) spatial patterns derived from a representative movie. For statistical comparison we then derived the mean of time-averaged stress for the analysed region. Average isotropic stress values correspond to subregion in bulk, away from boundaries.

### Two-photon laser ablation

Junctional tension of monolayers was examined by measuring the recoil of vertices after laser ablation of adherens junctions marked by E-Cadherin-mCherry cells, as previously described (Liang et al., 2016). A region of constant area (3.8 × 0.6 μm, with the longer axis orthogonal to the junctions) of the apical junctions was ablated by a femtosecond infrared two-photon 790 nm laser with 20% transmission and 15 iterations. The amount of recoil after ablation was measured as the change in the distance *L* between the vertices at *t* = *t* with respect to *t*=0 [i.e. *L*(*t*) − *L*(0)] of the ablated contacts as a function of time (*t*). At least 15 junctions were ablated and quantified for each condition of every experiment and the mean values of ε(*t*) were plotted against time to determine the extent of junctional initial recoil velocity after ablation using non-linear regression.

### RhoA FRET

FRET measurements and analysis were performed 48 hours after transfection. Imaging was performed on an LSM710 Zeiss confocal microscope. The donor channel was acquired using a 458 nm laser line and emission was collected in the donor emission region (470–490 nm). The acceptor channel was acquired using a 514 nm laser line and emission was collected in the acceptor emission range (530–590 nm). FRET signal was acquired using a 458 nm laser line and emission was collected in the acceptor emission range (530–590 nm). RhoA activation images were generated by calculating FRET/CFP emission ratios.

### FRAP

FRAP measurements and analysis were performed 36 hours after transfection with E-cadherin-mCherry. A ROI in the centre of cell-cell contacts was bleached using a 790 nm laser followed by 5 minutes of imaging with a frame interval of 1 second. To generate FRAP profiles, mean grey values at the bleached ROIs were measured during the recovery phase. The mean grey values obtained from the bleached region were normalized to the average pre-bleach value and fitted to a one phase exponential association function using GraphPad Prism software.

### Junctional protein quantification

Junctional intensity of E-cadherin, NMIIA, NMIIB, F-actin, α-catenin and α18 was quantified by generating line-scans of 20 μm lines perpendicular to randomly selected junctions. Line pixel intensities were then fitted with a Gaussian curve and peak values were obtained from the fitting. Pixel intensity values of 10 pixels on either side of the centres were considered background and subtracted from the peak values. Due to heterogenous expression levels of GFP-AHPH, the pixel values of 10 pixels on either side of the centers were divided from the peak values. A minimum of 60 junctions was measured in each experimental condition. The relative values of experimental conditions were normalised to the value of control cells for comparison.

### Statistical analysis

Statistical significance of the data was analysed by calculating the P-values using two-tailed students t-tests or one-way ANOVA test. Statistical analysis is shown as follows: n.s., not significant; *,P<0.05; **,P<0.01; ***,P<0.001; ****,P<0.0009.

### Code availability

The Matlab code for BISM tissue level tension analysis is available on request.

## Supplementary Figure captions

**Fig S1. Lamellipodia were evident throughout the motile population**

(A) Fluorescence images showing active lamellipodia expressing Lifeact TagRFPT at leading margin and rear edge of monolayer. Scale bar: 20μm.

**Fig S2. Active GTP-RhoA levels and localisation**

(A) Fluorescence images showing raw and intensity adjusted AHPH images at 0 min and 12 hours post migration.

(B) AHPH localisation in lamellipodia of Control and Anillin KD

(C) Change in junctional RhoA activation during MCF-7 monolayer migration measured using a FRET-based activity sensor. YFP and RhoA FRET (CFP/YFP) emission ratios images measured at ZA of cells during CCM. The graph shows an increase in RhoA FRET index after 12 hours of migration.

Data are means ± SEM; n=3 independent experiments. ****,P<0.0009; unpaired t-test. Scale bar: (B) 20μm (C) 10μm.

**Fig S3. Immunoblot images showing the confirmation of knockdown and rescue gene**

**Fig S4. Localisation of rescue construct at the ZA of MCF-7 cells and confirmation with Western blot**

## Notes

### Competing Interest Statement

The authors have declared no competing interest.

