## Supplementary figures and images for "Enhanced RhoA signaling stabilizes E-cadherin in migrating epithelial monolayers"

### Supplemental figures

A

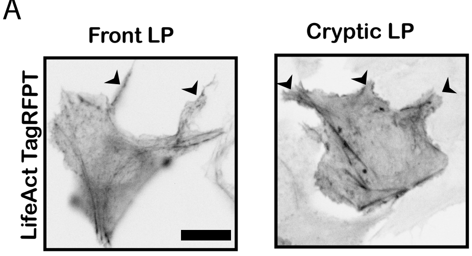


**Fig S1**

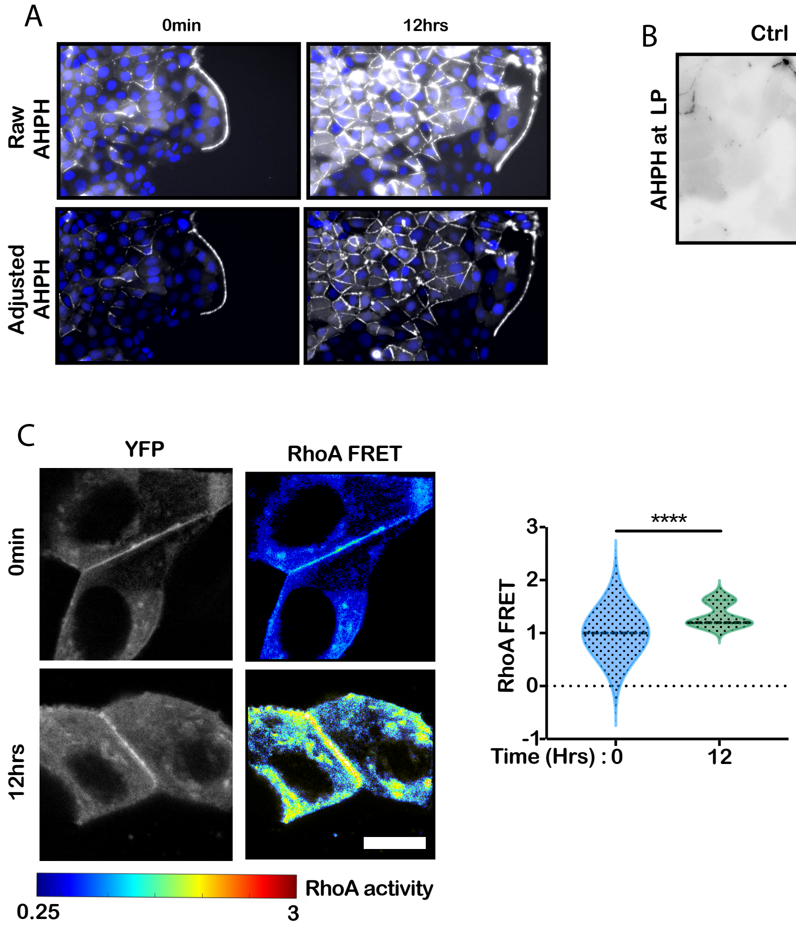

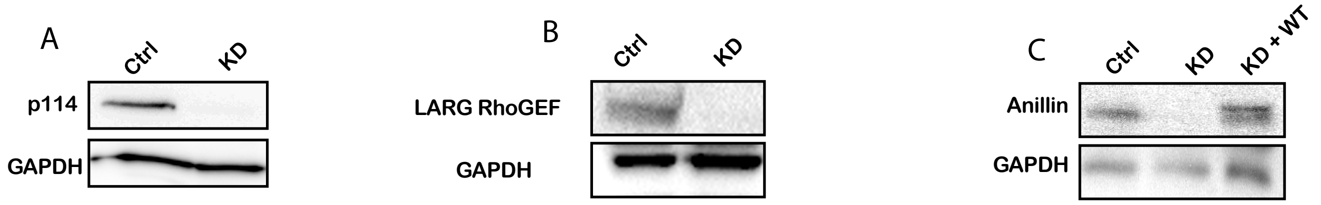

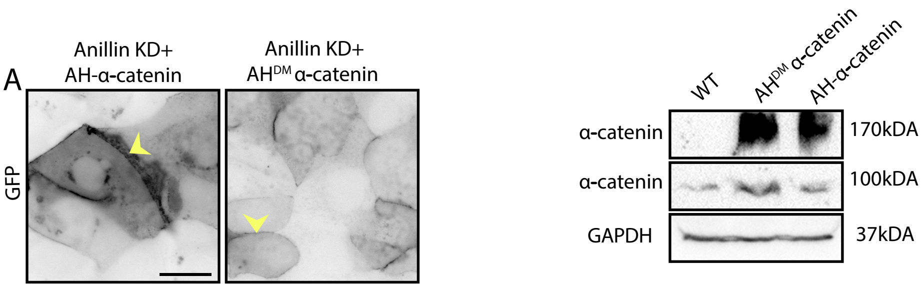
